# H3K18 & H3K23 acetylation directs establishment of MLL-mediated H3K4 methylation

**DOI:** 10.1101/2024.05.13.590588

**Authors:** Geoffrey C. Fox, Karl F. Poncha, B. Rutledge Smith, Lara N. van der Maas, Nathaniel N. Robbins, Bria Graham, Jill M. Dowen, Brian D. Strahl, Nicolas L. Young, Kanishk Jain

## Abstract

In an unmodified state, positively charged histone N-terminal tails engage nucleosomal DNA in a manner which restricts access to not only the underlying DNA, but also key tail residues subject to binding and/or modification. Charge-neutralizing modifications, such as histone acetylation, serve to disrupt this DNA-tail interaction, facilitating access to such residues. We previously showed that a polyacetylation-mediated chromatin “switch” governs the read-write capability of H3K4me3 by the MLL1 methyltransferase complex. Here, we discern the relative contributions of site-specific acetylation states along the H3 tail and extend our interrogation to other chromatin modifiers. We show that the contributions of H3 tail acetylation to H3K4 methylation by MLL1 are highly variable, with H3K18 and H3K23 acetylation exhibiting robust stimulatory effects, and that this extends to the related H3K4 methyltransferase complex, MLL4. We show that H3K4me1 and H3K4me3 are found preferentially co-enriched with H3 N-terminal tail proteoforms bearing dual H3K18 and H3K23 acetylation (H3{K18acK23ac}). We further show that this effect is specific to H3K4 methylation, while methyltransferases targeting other H3 tail residues (H3K9, H3K27, & H3K36), a methyltransferase targeting the nucleosome core (H3K79), and a kinase targeting a residue directly adjacent to H3K4 (H3T3) are insensitive to tail acetylation. Together, these findings indicate a unique and robust stimulation of H3K4 methylation by H3K18 and H3K23 acetylation and provide key insight into why H3K4 methylation is often associated with histone acetylation in the context of active gene expression.

## Introduction

Histone post-translational modifications (PTMs) engage in crosstalk, wherein a PTM(s) influences the deposition, removal, or recognition of another, thereby generating a combinatorial “histone code” [1]. Select combinations of PTMs direct distinct epigenetic landscapes through modulation of DNA accessibility and dictating which factors are recruited to or excluded from a given region of chromatin [2,3]. Though several key examples of histone PTM crosstalk have been elucidated, such as the stimulation of Gcn5’s histone acetyltransferase (HAT) activity by H3 serine 10 phosphorylation (H3S10ph) and H3 lysine 14 acetylation (H3K14ac), the relative lack of availability of recombinant, homogenously-modified nucleosomes – the fundamental repeating unit of chromatin – for *in vitro* interrogation of histone PTM crosstalk have hampered in-depth exploration of crosstalk mechanisms [4–6]. Biophysical characterization of the nucleosome has revealed that histone tails, which are enriched for positively charged residues, are often bound to the negatively charged nucleosomal DNA backbone, and therefore unable to be accessed by epigenetic machinery [7]. Acetylation of histone lysine residues, which neutralizes their charge, has recently been shown to release tails off of nucleosomal DNA in a manner which increases the accessibility of key residues for binding and/or modification by chromatin factors [8].

We previously showed that this form of PTM crosstalk is critical for the establishment of H3K4 methylation by the mixed-lineage leukemia 1 (MLL1/KMT2A) complex [9]. Making use of *in vitro* methylation assays with the recombinant MLL1 core complex in tandem with middle-down mass spectrometry for the quantification of histone PTMs, we found that a polyacetylation-mediated chromatin “switch” governs the read-write capability of MLL1 in the manifestation of higher-order H3K4 methylation (i.e., H3K4me3). This study provided the first biochemical insights into why MLL1-mediated H3K4me3 is so frequently associated with histone acetylation inside of cells, particularly at the promoters of actively transcribed genes [10].

Here, we expand on this work by showing that site-specific acetylation states differentially influence the activity of MLL1 *in vitro*, with a pronounced stimulation by H3K18ac and H3K23ac, and that this effect is shared with the related H3K4 monomethyltransferase, MLL4 (KMT2D). Middle-down mass spectrometry supports the preferential co-enrichment of H3K4me1 and H3K4me3 with H3 N-terminal tail proteoforms bearing the dual modification H3{K18acK23ac}. We show that this effect is unique to MLL family-mediated H3K4 methylation through the assessment of histone lysine methyltransferases targeting other H3 residues, both on the tail and on the nucleosome core, and the H3T3 kinase, Haspin. This work, making use of site-specific, homogenously-modified nucleosome substrates in tandem with middle-down mass spectrometry provides key insight into why H3K4 methylation specifically is so often associated with acetylation of active *cis*-regulatory elements, while other methylation events occurring elsewhere on histones, as well as a modification occurring directly adjacent to H3K4 are not [10,11].

## Results

### Introduction to combinatorial PTM/proteoform notation

This works focuses on crosstalk between histone post-translational modifications (PTMs) functioning in *cis*, on the same molecule. We use approaches that are capable of quantitively measuring single molecule co-occurrences of multiple PTMs and use reagents that contain defined sets of PTMs. The objective of this work is to understand the principles of enzymatic specificity and activity in the context of intact nucleosomes with combinatorial modifications present. To this end we make use of nomenclature, concepts, and notation to describe several related concepts concisely and unambiguously: Proteoforms, PTM combinations, and PTMs. A proteoform is the complete state of a protein, including defined occupancy of sites of variable modification [12]. Here we represent proteoforms as a sparse matrix by use of square brackets, e.g. H3[K18acK23ac], where unmodified sites are omitted for conciseness and clarity. We represent combinations of PTMs with curly brackets, e.g., H3{K18acK23ac}, where the state of other modification sites are undefined, variable, unknown, or ignored. For individual modifications, we use H3[K18ac] to indicate that it is the exclusive PTM present, and we use H3K18ac to indicate non-exclusive presence of this PTM. It is important to note that *in vivo* proteoforms are constrained. Minimal combinations of PTMs that are biochemically sufficient to modulate enzymatic activity *in vitro* often occur exclusively with additional PTMs that have little direct effect but are prerequisite. Our focus here is the direct effect on enzymatic activity but we also note the *in vivo* prerequisites.

### Acetylation of H3K18 & H3K23 stimulates MLL4-mediated H3K4 methylation in vitro and in vivo

We first sought to understand the contribution of site-specific H3 acetylation (H3ac) states to H3K4 methylation by performing *in vitro* methylation assays with the recombinant MLL4 core complex (KMT2D; MLL4^SET^/WDR5/RbBP5/ASH2L/DPY30), responsible for catalyzing H3K4 monomethylation (H3K4me1) *in vivo*, and a panel of differentially monoacetylated nucleosome substrates (**Figure 1A**) [13]. Endpoint analysis revealed that while the contribution of H3ac to MLL4-mediated H3K4 methylation is highly variable, H3K18ac, H3K23ac, and H3K36ac have a pronounced stimulatory effect relative to unmodified nucleosomes. H3K9ac and H3K14ac displayed a more modest, yet reproducible stimulatory effect. Notably, H3K27ac – a PTM frequently associated with H3K4me1 at active enhancers – had no effect on MLL4-mediated H3K4 methylation *in vitro* [14–16]. Surprisingly, the proximity of the acetylated residue to H3K4 is not a primary determinant of the magnitude of the observed stimulation. Rather, acetylation events occurring toward the center of the H3 N-terminal tail (residues 9-23), and those occurring toward the nucleosome core (H3K36) have the most pronounced effect.

**Figure 1.**
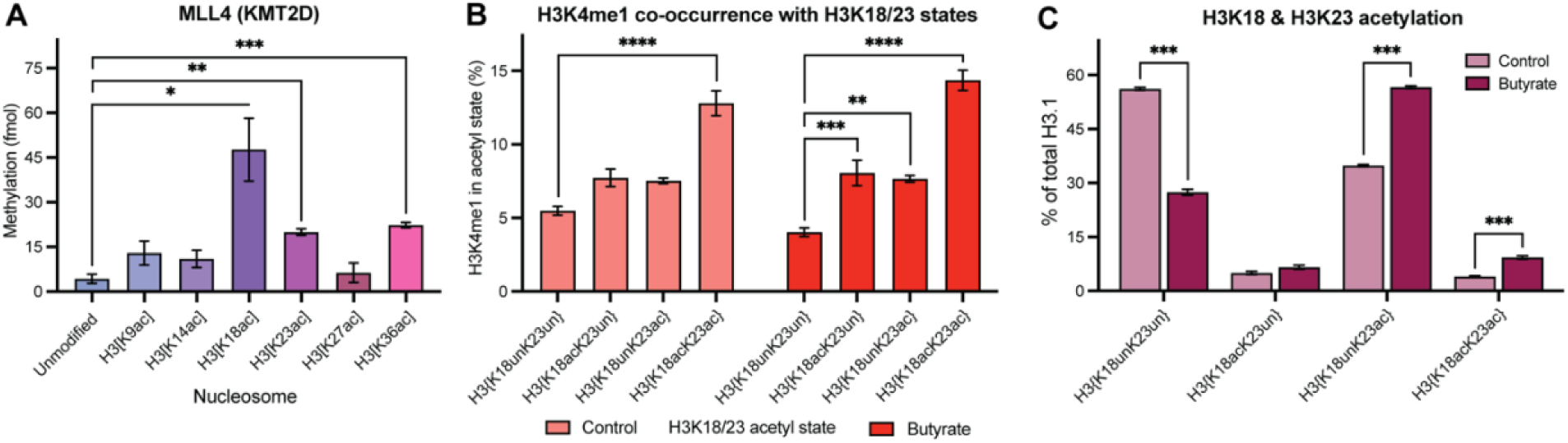
Acetylation of H3K18 & H3K23 promotes H3K4me1. (A) *In vitro* methylation assays with the recombinant MLL4 complex (10 nM) and a panel of designated nucleosomes (300 nM) reveal a robust stimulation of MLL4-mediated H3K4 methylation by H3K18ac & H3K23ac. (B) Middle-down MS of HEK293 cells quantifying co-enrichment of H3K4me1 with distinct H3K18/23 acetylation states with (dark red) and without (light red) histone deacetylase inhibition (butyrate). Changes in site-specific N-terminal H3 tail acetylation states as a result of butyrate treatment are shown in Supplementary Figure S1. (C) Quantification of total H3K18 and H3K23 acetylation states by middle-down mass spectrometry. MS data shown is reported in Supplementary Table 1. Significance was determined by Student’s t-test. NS unless otherwise designated. \**p* < 0.05, \*\**p* < 0.005, \*\*\**p* < 0.001, \*\*\*\**p* < 0.0001. *n* = 3. Error: SEM.

We then sought to understand the contribution of site-specific H3 acetylation to MLL4-mediated H3K4me1 *in vivo* by analyzing our existing middle-down mass spectrometry dataset from HEK293 cells with and without the inhibition of histone deacetylases (sodium butyrate; HDACi) to modulate the acetylation landscape (**Figure S1**) [9]. We find that H3K4me1 is preferentially enriched on H3 N-terminal tails bearing dual H3{K18acK23ac} acetylation (**Figure 1B**). Notably, H3K4me1 is only modestly enriched with H3 N-terminal tail proteoforms bearing single H3K18 or H3K23 acetylation relative to H3 tail proteoforms bearing the dual acetylation, H3{K18acK23ac}. This suggests that H3 tail proteoforms bearing multiple acetylated residues are a preferable substrate for MLL4-mediated H3K4 methylation *in vivo*, with a synergistic contribution from H3K18 and H3K23 acetylation specifically. Moreover, under HDACi, H3K4me1 co-enrichment with either H3K18ac- or H3K23ac-bearing proteoforms becomes significant. The global levels of H3K4me1 remain largely unchanged under 2-hours of HDACi (**Figure S2A**). This is consistent with our previous work suggesting that acetylation more tightly governs the deposition of higher-order H3K4 methyl states (i.e., MLL1-mediated H3K4me3) *in vivo*, as well as previous work displaying hours-scale stability of global H3K4me1 levels upon even direct perturbation of its deposition [9,17]. However, it is noteworthy that HDACi raises the global levels of H3{K18acK23ac} proteoforms approximately 2.3-fold (*p* = 0.00077), and the co-occurrence of H3K4me1 with these proteoforms remains consistent, reflecting a proportional gain in H3K4me1 on the same H3{K18acK23ac} tail that gains acetylation (**Figures 1B & 1C**). Further mechanistic studies of how acetylation of H3K18 and/or H3K23 modulate N-terminal H3 tail conformation and dynamics could elucidate how these events may be uniquely poised to govern MLL-mediated H3K4 methylation.

### The stimulatory effect of H3K18ac & H3K23ac extends to MLL1-mediated H3K4 methylation in vitro and in vivo

Having explored the site-specific contributions of H3 tail acetylation to MLL4-mediated H3K4 methylation, we asked if these stimulation patterns observed with MLL4 extended to the related H3K4 methyltransferase, MLL1 (KMT2A), responsible for catalyzing H3K4 trimethylation (H3K4me3) *in vivo*. We performed *in vitro* methylation assays with the recombinant MLL1 core complex (MLL1^SET^/WDR5/RbBP5/ASH2L/DPY30) and the same panel of differentially monoacetylated nucleosomes. We observe stimulation of MLL1-mediated H3K4 methylation that resembled that which was observed with MLL4 (**Figure 2A**) [18]. Specifically, a similarly robust increase in MLL1-mediated H3K4 methylation is observed on nucleosomes with H3K18ac, and to a lesser extent, H3K23ac. H3K9ac, H3K14ac, and H3K36ac also modestly stimulated MLL1 activity. H3K27ac again has no impact on MLL1-mediated H3K4 methylation, another direct parallel to that which was observed with MLL4.

**Figure 2.**
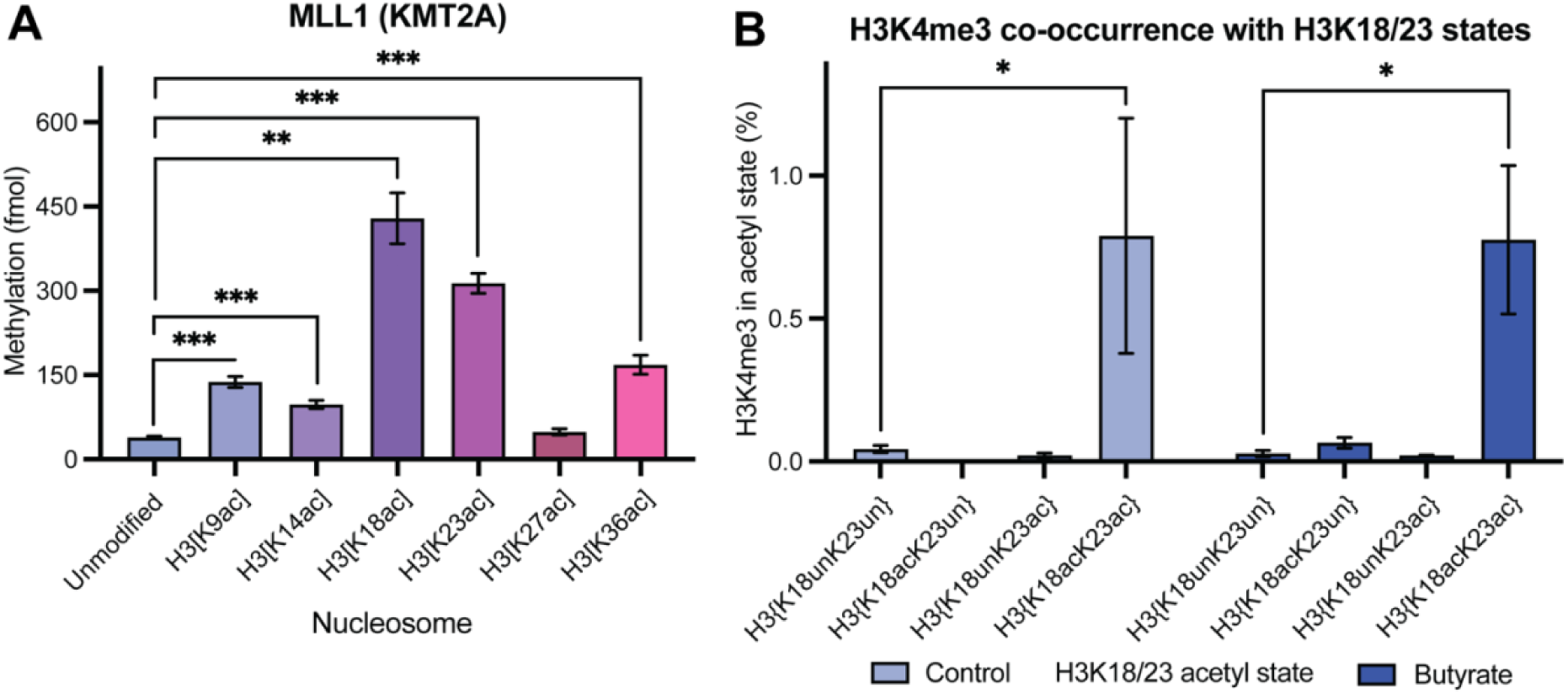
Acetylation of H3K18 & H3K23 promotes H3K4me3. (A) *In vitro* methylation assays with recombinant MLL1 (10 nM) and a panel of designated nucleosomes (300 nM) reveal a robust stimulation of MLL1-mediated H3K4 methylation by H3K18ac & H3K23ac. (B) Middle-down MS of HEK293 cells quantifying co-enrichment of H3K4me3 with distinct H3K18/23 acetylation states with (dark blue) and without (light blue) histone deacetylase inhibition (butyrate). MS data shown is reported in Supplementary Table 1. Significance was determined by Student’s t-test. NS unless otherwise designated. \**p* < 0.05, \*\**p* < 0.005, \*\*\**p* < 0.001. *n* = 3. Error: SEM.

Our previous studies of how H3 acetylation contributes to H3K4me3 *in vivo* revealed that an H3 polyacetylation-mediated chromatin “switch” governs H3K4me3 read-write capability, whereby only proteoforms bearing H3 polyacetylation are significantly enriched for H3K4me3 [9]. We therefore asked if H3 proteoforms enriched for H3K18ac and H3K23ac were found specifically co-enriched for H3K4me3 by analyzing our existing middle-down mass spectrometry dataset from HEK293 cells with and without HDACi. As expected, there is a strong preferential co-enrichment for H3K4me3 on H3 proteoforms bearing the dual acetyl mark H3{K18acK23ac} relative to proteoforms bearing either H3K18ac or H3K23ac alone and proteoforms bearing neither (**Figure 2B**). In fact, very little H3K4me3 is detected on proteoforms bearing single or no acetylation at H3K18 and H3K23, which holds true under HDACi, contrary to findings from our experiments with MLL4 & H3K4me1. This falls in-line with our previously reported model of how polyacetylation (i.e., tetra- and penta-acetylation) of the H3 tail robustly enhances MLL1-mediated H3K4me3 *in vivo* [9]. Indeed, 2-hour HDACi treatment – which significantly increases the presence of polyacetylated H3 proteoforms – increases total H3K4me3 levels approximately 2.2-fold (**Figure S2A**). Like MLL4-mediated H3K4me1, the co-occurrence of MLL1-mediated H3K4me3 with H3{K18acK23ac} proteoforms is consistent under HDACi conditions where the abundance of H3{K18acK23ac} is increased approximately 2.3-fold, again reflecting a proportional gain in H3K4me3 on the same tail as H3{K18acK23ac} as that tail gains acetylation (**Figures 2B & 1C**). Though the MLL1 core complex can methylate monoacetylated substrates *in vitro* with relative efficiency, it is clear that the process is more tightly regulated inside of the nucleus. Further *in vivo* studies identifying additional co-regulators of MLL1-mediated H3K4me3 contributing to this discrepancy are warranted.

### Methyltransferases targeting other H3 tail lysine residues are insensitive to H3ac in vitro

To understand whether the stimulatory effect of H3ac extended to methyltransferases targeting other H3 N-terminal tail lysine residues, we performed *in vitro* methylation assays with G9a (H3K9me1/2 writer *in vivo*), SETDB1 (H3K9me3 writer *in vivo*), the polycomb repressive complex 2 (PRC2; H3K27me writer *in vivo*), and ASH1L (H3K36me2 writer *in vivo*) and panels of differentially monoacetylated nucleosomes (**Figure 3A-D**) [19–22]. In general, little to no H3 acetylation-mediated enhancement of H3 methylation on the appropriate lysine residue is observed when testing non-H3K4 methylation of histone H3 on nucleosomes. Certain statistically significant, if modest, effects were observed, however; G9a is modestly (1.2-fold; *p* = 0.0399) stimulated by H3K18 acetylation, while the only other significant contributions observed by the distinct acetylation states were inhibitory in the cases of H3K27ac and H3K36ac (**Figure 3A**). Similarly, the activity of SETDB1 is inhibited by H3K36ac (**Figure 3B**). In contrast to the H3K4 methyltransferases, G9a, SETDB1, PRC2, and ASH1L exhibit robust activity against unmodified nucleosome substrates, suggesting that methylation carried out by these enzymes is a process less tightly governed by acetylation relative to H3K4 methylation.

**Figure 3.**
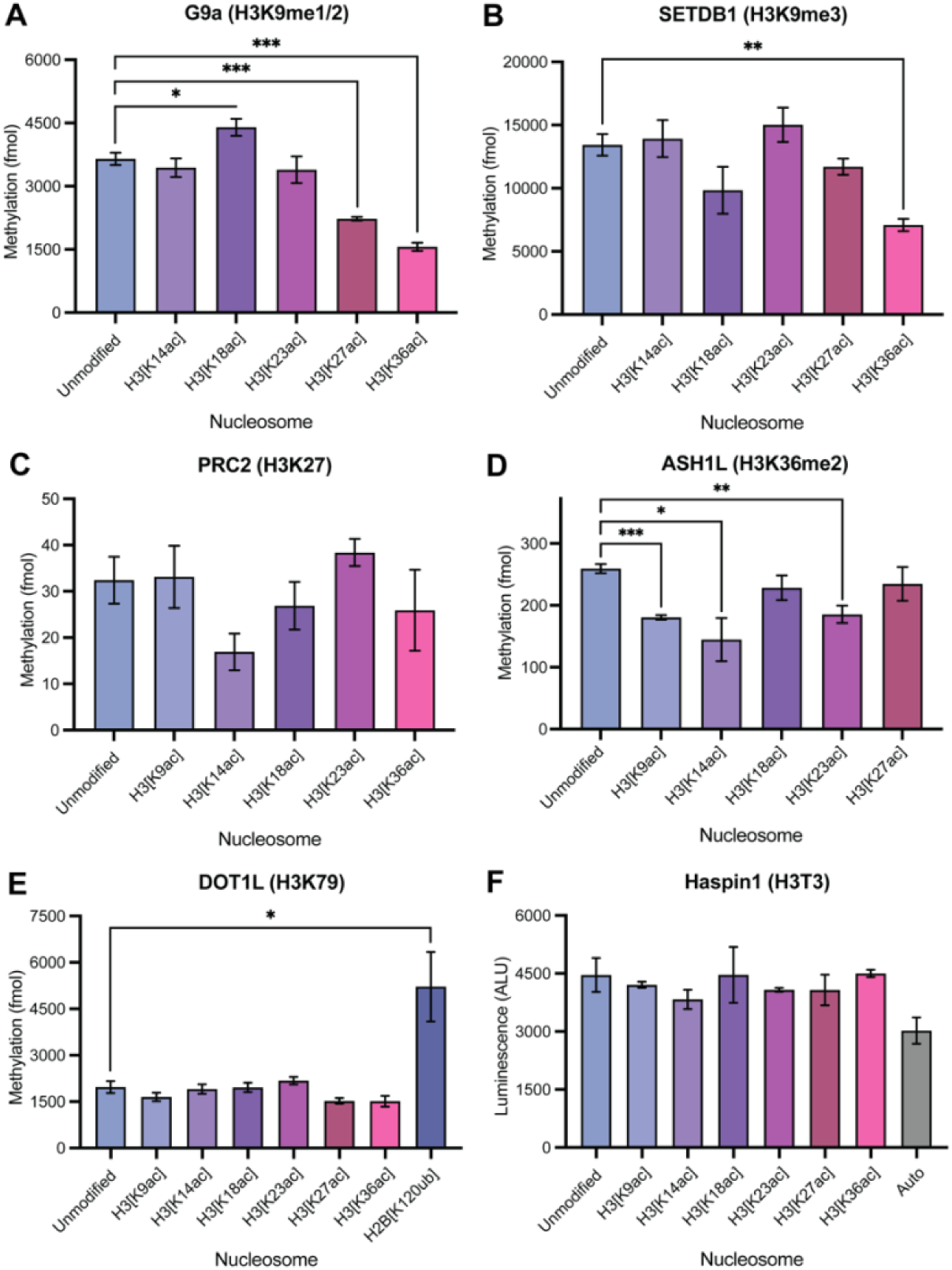
Methyltransferases performing non-H3K4 H3 methylation are insensitive to acetylation. (A-E) *In vitro* methylation assays with (A) recombinant G9a (913-1193; 10 nM), (B) recombinant SETDB1 (567-1291; 25 nM), (C) recombinant PRC2 (EED/EZH2/SUZ12/AEBP/RbAp48; 50 nM), (D) recombinant ASH1L (2046-2330; 50 nM), or (E) recombinant DOT1L (1-420; 15 nM) and a panel of designated nucleosomes (300 nM) reveal minimal influence of H3 acetylation on non-H3K4 lysine methylation. (F) *In vitro* kinase assays with recombinant Haspin (465-798; 25 nM) and a panel of designated nucleosomes (*EpiCypher*) reveal minimal influence of H3 acetylation state on Haspin-mediated H3T3 phosphorylation. Auto: autophosphorylation (no substrate). Significance was determined by Student’s t-test. NS unless otherwise designated. \**p* < 0.05, \*\**p* < 0.005, \*\*\**p* < 0.001. *n* = 3. Error: SEM.

To assess the degree to which acetylation state influenced non-H3K4 methylation of H3 lysine residues *in vivo*, we analyzed our existing middle-down mass spectrometry dataset from HEK293 cells with and without HDACi and quantified the global levels of H3K9, H3K27, and H3K36 methylation states (**Figures S2B-D & S3**). While HDACi manifested a modest shift in global H3K9 methylation states to H3K9me2 and H3K9me3, total levels of H3K9 methylation remain largely unchanged under HDACi. Indeed, like H3K9 methylation, H3K27 methylation and H3K36me2 are also unaffected by HDACi (**Figures S2C-D & S3**).

To determine whether the influence of H3 acetylation-mediated accessibility is limited to methyltransferases targeting the H3 N-terminal tail itself, we performed *in vitro* methylation assays with recombinant DOT1L, a methyltransferase targeting a residue embedded within the nucleosome core, H3K79 – considerably spatially segregated from the H3 tail – and the same panel of differentially monoacetylated nucleosomes (**Figure 3E**) [23]. As expected, DOT1L is insensitive to N-terminal H3 tail acetylation and exhibits robust activity against the nucleosome regardless of the H3ac state. As a positive control, we also tested DOT1L activity against nucleosome substrates bearing ubiquitylation of H2B at lysine 120 (H2B[K120ub]), a PTM previously reported to enhance the deposition of DOT1L-mediated H3K79 methylation [24,25]. As expected, DOT1L activity is significantly stimulated by H2BK120ub. These results suggest that acetylation of the histone tails does not drastically influence the conformation of the nucleosome core in a way that facilitates access to modifiable residues within it, such as H3K79. Rather, the impact of histone tail acetylation on the accessibility of other modifiable histone residues may instead be primarily restricted to that same tail.

### Haspin-mediated H3T3 phosphorylation is insensitive to H3ac

Last, in order to determine if the enhanced accessibility to H3K4 by H3K18ac and H3K23ac is generalizable to the residues present at the N-terminal end of the H3 tail, or if it is specific to H3K4, we performed *in vitro* kinase assays with Haspin, a kinase targeting H3 threonine 3 (H3T3) and the same panel of differentially acetylated nucleosomes (**Figure 3F**) [26].

Surprisingly, neither H3K18ac nor H3K23ac stimulated the deposition of H3T3ph by Haspin. In fact, none of the assessed acetylation events significantly stimulated Haspin-mediated H3T3ph. This suggests that H3K18ac and H3K23ac may serve to selectively poise H3K4 for modification, perhaps only by MLL family methyltransferases, but not other neighboring residues present at the H3 N-terminus.

## Discussion

Our understanding of the complexity of the histone code has grown over the last twenty years, thanks in part to cutting edge mass spectrometry and genomic approaches. Numerous studies have amassed compelling correlative data that link certain histone PTMs to each other, such as histone H3 acetylation and H3K4 methylation [27–31]. On top of this, recent structural and biochemical analysis of histone PTMs on nucleosomes rather than peptides have revealed new mechanisms for regulating these biologically observed correlations (i.e., co-occurrence of H3K4 methylation and H3 acetylation on regions of actively transcribing genes) [7–9]. As technologies improve our ability to more discretely and quantitatively study the complexity of histone PTM crosstalk that occurs biologically, we have followed up our initial discovery of polyacetylation-mediated activation of H3K4 methylation through further use of modified nucleosomes, enzymology, and quantitative mass spectrometry.

By pairing *in vitro* methylation and kinase assays with middle-down mass spectrometry for quantifying histone PTMs, we report a unique and robust governance of H3K4 methylation by H3K18 and H3K23 acetylation (**Figures 1 & 2**). H3K4 methyltransferases MLL1 (KMT2A) & MLL4 (KMT2D) are each sensitive to site-specific acetylation of either of these residues *in vitro* (**Figures 1A & 2A**). Moreover, the similarities in the sensitivities of MLL1 & MLL4 to site-specific H3 tail acetylation extend to other residues which stimulate their activities *in vitro*, such as H3K9ac & H3K36ac, and even those which don’t, like H3K27ac. This suggests that similar mechanisms of acetylation-mediated H3K4 accessibility may apply across enzymes that require such access.

Inside of cells, there is a marked preferential co-enrichment of H3K4me1 and H3K4me3 with H3 proteoforms bearing dual H3{K18acK23ac} in *cis* relative to both mono- and unacetylated proteoforms (**Figures 1B & 2B**). This preference is particularly apparent with H3K4me3, which can hardly be detected on proteoforms lacking dual H3{K18acK23ac}. We also observed an increase in global H3K4me3 under HDACi (**Figure S2A**). This is consistent with our previous studies of MLL1, where we found that H3K4me3 deposition is only permitted on H3 tails also bearing polyacetylation [9].

Finally, by analyzing other methyltransferase events along the H3 tail such as G9a, SETDB1, PRC2, and ASH1L, and Haspin, a kinase that phosphorylates H3T3, we showed that these non-H3K4 modifying enzymes were not affected by H3 N-terminal tail acetylation. These results were largely expected, as G9a, SETDB1, and the PRC2 complex deposit histone PTMs (H3K9 and H3K27 methylation) that are fundamentally associated with repression of gene expression and at odds with the acetylation-mediated opening of chromatin. Notably, our results contrast findings indicating a stimulation of G9a-mediated H3K9 methylation as a result of the enzymatic acetylation of recombinant nucleosomes [32]. We do not observe a similar stimulation of G9a on our panel of homogeneously monoacetylated recombinant substrates, nor can we detect a significant impact of the elevated acetylation landscape manifested by HDACi on global H3K9 methylation by middle-down MS (**Figure S3**). Somewhat surprisingly, ASH1L, a writer for H3K36me2 – a histone PTM associated with transcription elongation – is also largely unaffected by histone H3 acetylation on nucleosomes, supported by the lack of increase in H3K36 methylation as a function of HDACi *in vivo* (**Figure S2**). The lack of stimulation of Haspin-mediated H3T3 phosphorylation by H3 acetylation further highlighted that sensitivity of MLL-mediated H3K4 methylation to H3K18ac and H3K23ac is not simply a regional effect extending across the residues present at the extreme N-terminal end of H3, but rather, is more specifically targeted to H3K4 methylation.

Taken together, these findings highlight a unique and robust governance of H3K4 accessibility by H3K18ac and H3K23ac which extends neither to its nearest modifiable neighbor residue in H3T3, nor to other lysine residues along the H3 tail which are also subject to methylation (**Figure 4**). These findings give key insight into why H3K4 methylation specifically is consistently associated with histone acetylation *in vivo*, frequently found co-enriched at active *cis*-regulatory elements [10,11]. H3K18ac and H3K23ac are each deposited by the promiscuous acetyltransferase p300/CBP, which has previously been shown to be recruited to chromatin by MLL family complexes *in vivo* [33–35]. It is therefore feasible that MLL family H3K4 methyltransferases partake in a feedforward loop with p300/CBP, whereby p300/CBP-mediated acetylation promotes the deposition of H3K4 methylation by MLL, which recruits additional p300/CBP and MLL complexes (through its PHD reader domains) to ensure the robust enrichment of nucleosomes bearing both H3K4 methylation and acetylation that is capable of overwhelming the repressive regulatory machinery these factors share space within the nucleus with. Mechanistic studies characterizing how acetylation of H3K18 and/or H3K23 poise H3K4 for modification by MLL family H3K4 methyltransferases are warranted While some chromatin modifiers are effectively licensed by preexisting PTMs others are largely unaffected by the same PTMs. Interdependencies of this nature enable the tight, yet dynamic regulation of gene expression necessary to prevent spurious transcriptional activation and/or repression, and in turn, to promote appropriate transcriptional programs across cellular contexts. Thus, additional focus will be required on the interrogation of whether acetylation-mediated accessibility governs the activity of chromatin modifiers performing chemistry unique to that which is analyzed in this study (i.e., lysine demethylases, arginine methyltransferases, etc.), further building a foundation of knowledge for understanding how histone PTMs can function in combinatorial “codes” to govern genome function.

**Figure 4.**
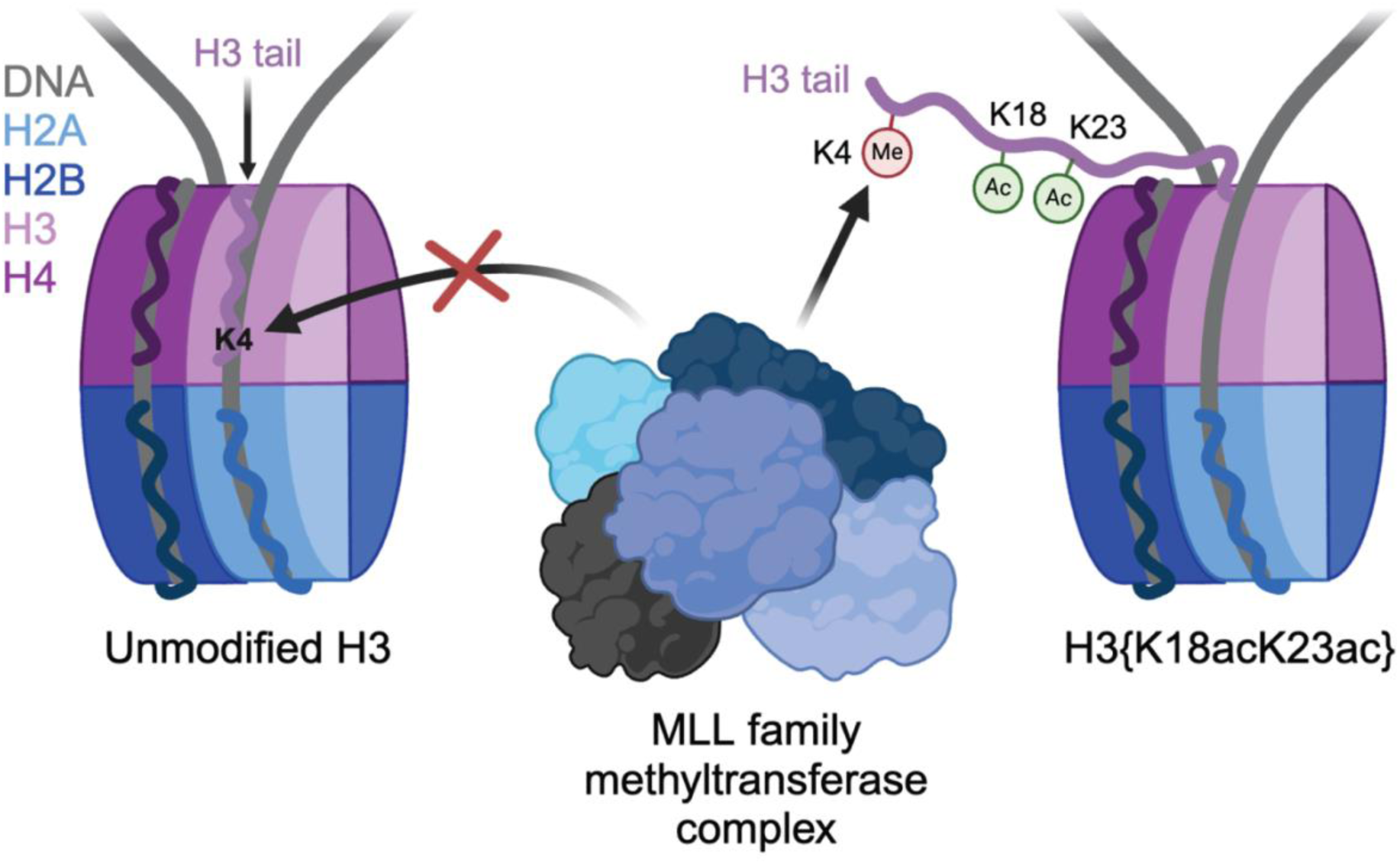
Model for H3{K18acK23ac}-mediated stimulation of MLL-mediated H3K4 methylation. In the unmodified state (left), the H3 N-terminal tail preferentially engages nucleosomal DNA in a manner that restricts access to H3K4 by MLL family methyltransferases. Acetylation of H3K18 and H3K23 (right) releases the H3 tail from nucleosomal DNA, permitting H3K4 access by MLL family methyltransferases.

## Experimental Procedures

### Expression, purification, and assembly of MLL/KMT2 family complexes

Methods for the expression, purification, and assembly of MLL4 core complex (MLL1 SET domain, WDR5, RbBP5, Ash2L and DPY30) were adapted from published protocols [36]. The MLL1 SET domain portion of a polycistronic recombinant expression construct containing the MLL1 SET domain (Uniprot Q03164; residues 3745-3969), WDR5 (Uniprot P61964; residues 2-334), RbBP5 (Uniprot Q15291; residues 1-538) and Ash2L (Uniprot Q9UBL3-3; residues 1-534) in pST44 vector (a kind gift from Dr. Song Tan) was swapped with the SET domain of MLL4 (Uniprot O14686; residues 5319-5538) and purchased from VectorBuilder [37]. The MLL4 core complex was purified identically to the MLL1 core complex, whose details can be found in our previous study [9].

### Plasmid construction

A recombinant expression construct encoding the G9a SET domain (Uniprot Q96KQ7; residues 913-1193) was received as a kind gift from Dr. Samantha Pattenden. A recombinant expression construct encoding the DOT1L catalytic domain (Uniprot Q8TEK3; residues 1-420) was purchased from AddGene (AddGene #36196). A recombinant expression construct encoding the Haspin kinase domain (GSG2; Uniprot Q8TF76; residues 465-798) was purchased from AddGene (AddGene #38915).

### Expression and purification of polyhistidine-tagged methyltransferase enzymes

Plasmids were transformed into chemically competent BL21.DE3(pLysS) *E. coli*, plated onto LB-agar plates containing the appropriate antibiotic (ampicillin or kanamycin), and incubated overnight at 37° C. One colony was inoculated into a LB preculture (containing either 50 µg/mL carbenicillin or 50 µg/mL kanamycin), incubated at 16° C overnight, transferred into 1 L LB containing the appropriate antibiotic, and cultured at 37° C until OD_600_ = 0.5∼0.7. Cultures were then transferred to 16° C and induced with 1 mM IPTG (*Sigma*) overnight. Following induction, cultures were harvested by centrifugation, and cell pellets were flash frozen in liquid nitrogen and stored at -20° C until use.

Cell pellets were resuspended in 50 mL of lysis buffer (25 mM sodium phosphate pH 7.4, 300 mM NaCl, 1 mM DTT, 1 mM PMSF, 20 mM imidazole supplemented with 1 mg/mL lysozyme (*Sigma*) and 250 U of Pierce Universal Nuclease (*ThermoFisher*)) and incubated at 37° C for 10 minutes. Cells were then lysed by sonication (5 x 30 seconds, 40% cycle, 40% power) and the lysate clarified by centrifugation. All chromatography was performed at 4° C. Clarified lysate was applied to a 5 mL HisTrap FF Ni-NTA column (*Cytiva*) equilibrated in IMAC wash buffer (25 mM sodium phosphate pH 7.4, 300 mM NaCl, 1 mM DTT, 1 mM PMSF, 20 mM imidazole) at 0.5 mL/min. Column was then washed with 20 column volumes (100 mL) of IMAC wash buffer. Bound polyhistidine-tagged proteins were then subjected to linear gradient elution from IMAC wash buffer to IMAC elution buffer (wash buffer supplemented with 250 mM imidazole) across 12 column volumes. 2 mL elution fractions were collected and assessed for purity by Coomassie-stained SDS-PAGE. Fractions containing pure polyhistidine-tagged proteins were pooled and concentrated by centrifugation filtration (*EMD Millipore*; using appropriate MWCO, either 10 or 30 kDa) and buffer exchanged into storage buffer (50 mM HEPES, pH 8.0, 150 mM NaCl, 1 mM DTT, 20% glycerol). Concentrated pools were aliquoted, quantified by Coomassie-stained SDS-PAGE-based densitometry with known BSA standards, and stored at -80° C until use.

### Expression and purification of polyhistidine-tagged Haspin

Plasmid was transformed into chemically competent BL21.DE3(pLysS) *E. coli*, plated onto LB-agar plates containing 50 µg/mL kanamycin, and incubated overnight at 37° C. One colony was inoculated into a 100 mL LB preculture (containing 50 µg/mL kanamycin), incubated at 16° C overnight, transferred into 1 L LB (containing 50 µg/mL kanamycin), and cultured at 37° C until OD_600_ = 0.5∼0.7. Cultures were transferred to 16° C and induced with 1 mM IPTG (*Sigma*) overnight. Following induction, cultures were harvested by centrifugation. Cell pellet was resuspended in 50 mL lysis buffer (25 mM sodium phosphate pH 7.4, 300 mM NaCl, 1 mM DTT, 1 mM PMSF, 20 mM imidazole), flash frozen in liquid nitrogen and stored at -20° C until use.

Frozen cell suspension was thawed at 37° C for approximately 15 minutes, supplemented with 1 mg/mL lysozyme (*Sigma*) and 250 U of Pierce Universal Nuclease (*ThermoFisher*), and incubated at 37° C for 10 minutes. Cells were then lysed by sonication (5 x 30 seconds, 40% cycle, 40% power) and the lysate clarified by centrifugation. All chromatography was performed at 4° C. Clarified lysate was applied to a 5 mL HisTrap FF Ni-NTA column (*Cytiva*) equilibrated in IMAC wash buffer (25 mM sodium phosphate pH 7.4, 300 mM NaCl, 1 mM DTT, 20 mM imidazole) at 0.5 mL/min. Column was then washed with 20 column volumes (100 mL) of IMAC wash buffer. Bound polyhistidine-tagged proteins were then subjected to linear gradient elution from IMAC wash buffer to IMAC elution buffer (IMAC wash buffer supplemented with 250 mM imidazole) across 12 column volumes. 2 mL elution fractions were collected and assessed for purity by Coomassie-stained SDS-PAGE. Fractions containing pure polyhistidine-tagged proteins were pooled and concentrated approximately 2-fold as described above. Concentrated pool (approx. 8 mL) was applied to a HiLoad Superdex 16/600 (120 mL) 75 pg preparative gel filtration column (*Cytiva*) equilibrated in GF buffer (50 mM Tris-Cl pH 8.0, 150 mM NaCl, 1 mM DTT) at 0.5 mL/min in four separate runs. 1.5 column volumes of GF buffer was used for elution and 2 mL peak fractions were collected, assessed for purity by Coomassie-stained SDS-PAGE, pooled according to presence of purified Haspin, and concentrated as described above. Glycerol was then added to a final concentration of 20%. Concentrated pools were aliquoted, quantified by Coomassie-stained SDS-PAGE-based densitometry with known BSA standards, and stored at -80° C until use.

### *In vitro* methylation assays (MLL1 and MLL4)

Methylation assays were performed as previously described with the following modifications [9]. For endpoint analysis, purified MLL1 (10 nM) or MLL4 (10 nM) was incubated with nucleosome substrate (300 nM) for 3 hours at 15° C following addition of 10 µM 9:1 *S*-adenosyl-L-methionine (SAM) *p*-toluenesulfonate salt (*Sigma*) to *S*-adenosyl-L-[*methyl*-^3^H]-methionine ([*methyl*-^3^H]-SAM) (*PerkinElmer*) in a reaction volume of 20 µL (in 50 mM HEPES, pH 8.0, 1 mM DTT, 1 µM ZnCl_2_). Following incubation, reactions were quenched by the addition of 5 µL of a 5X SDS loading dye. Reactions were separated by 15% Tris-Glycine SDS-PAGE, visualized by Coomassie staining and bands corresponding to histone proteins were excised. The excised bands were solubilized in a mixture of 50% Solvable (*PerkinElmer*) and 50% water for 3 hours at 50° C. Mixture and gel slices were transferred to scintillation vials and 10 mL Hionic-Fluor scintillation fluid (*PerkinElmer*) was added, vortexed briefly, dark-adapted overnight, and measured for radioactivity on a Liquid Scintillation Counter (*Beckman-Coulter*).

### *In vitro* methylation assays (G9a and SETDB1)

Methylation assays were performed as previously described with the following modifications [38]. Optimal enzyme concentrations for endpoint analysis were identified through an enzyme titration (0-300 nM for G9a, 0-200 nM for SETDB1) against 1 µg of chicken erythrocyte oligonucleosomes according to the procedure described below (**Figure S4A-B**). Purified 6xHIS-G9a (913-1193; 10 nM) or GST-SETDB1 (567-1291; 25 nM; a kind gift from Dr. Samantha Pattenden) was incubated with nucleosome substrate (300 nM) for 1 hour at 30° C following addition of 10 µM 9:1 *S*-adenosyl-L-methionine (SAM) *p*-toluenesulfonate salt (*Sigma*) to *S*-adenosyl-L-[*methyl*-^3^H]-methionine ([*methyl*-^3^H]-SAM) (*PerkinElmer*) in a reaction volume of 20 µL (in 50 mM Tris-Cl, pH 8.8, 5 mM MgCl_2,_ 4 mM DTT). Downstream assessment of methylation was performed as described with MLL methylation assays.

### *In vitro* methylation assays (PRC2)

Methylation assays were performed as previously described with the following modifications [39]. Optimal enzyme concentration for endpoint analysis was identified through an enzyme titration (0-100 nM) against 1 µg of chicken erythrocyte oligonucleosomes according to the procedure described below (**Figure S4C**). Purified PRC2 (EED/EZH2/SUZ12/AEBP/RbAp48; 50 nM; *BPS Biosciences*) was incubated with nucleosome substrate (300 nM) for 1 hour at 30° C following addition of 10 µM 9:1 *S*-adenosyl-L-methionine (SAM) *p*-toluenesulfonate salt (*Sigma*) to *S*-adenosyl-L-[*methyl*-^3^H]-methionine ([*methyl*-^3^H]-SAM) (*PerkinElmer*) in a reaction volume of 20 µL (in 50 mM Tris-Cl, pH 8.5, 5 mM MgCl_2,_ 4 mM DTT). Downstream assessment of methylation was performed as described with MLL methylation assays.

### *In vitro* methylation assays (ASH1L)

Optimal enzyme concentration for endpoint analysis was identified through an enzyme titration (0-200 nM) against 1 µg of chicken erythrocyte oligonucleosomes according to the procedure described below (**Figure S4D**). Purified ASH1L (2046-2330; 50 nM; *Reaction Biology*) was incubated with nucleosome substrate (300 nM) for 1 hour at 30° C following addition of 10 µM 9:1 *S*-adenosyl-L-methionine (SAM) *p*-toluenesulfonate salt (*Sigma*) to *S*-adenosyl-L-[*methyl*-^3^H]-methionine ([*methyl*-^3^H]-SAM) (*PerkinElmer*) in a reaction volume of 20 µL (in 50 mM Tris-Cl pH 9.0, 1.5 mM MgCl_2_, 1 mM DTT). Downstream assessment of methylation was performed as described with MLL methylation assays.

### *In vitro* methylation assays (DOT1L)

Methylation assays were performed as previously described with the following modifications [40]. Optimal enzyme concentration for endpoint analysis was identified through an enzyme titration (0-150 nM) against 1 µg of chicken erythrocyte oligonucleosomes according to the procedure described below (**Figure S4E**). Purified DOT1L (1-420; 15 nM) was incubated with nucleosome substrate (300 nM) for 30 minutes at 30° C following addition of 10 µM 9:1 *S*-adenosyl-L-methionine (SAM) *p*-toluenesulfonate salt (*Sigma*) to *S*-adenosyl-L-[*methyl*-^3^H]-methionine ([*methyl*-^3^H]-SAM) (*PerkinElmer*) in a reaction volume of 20 µL (in 20 mM Tris-Cl pH 8.0, 50 mM NaCl, 1 mM DTT, 1 mM EDTA, 0.1 mg/mL BSA). Downstream assessment of methylation was performed as described with MLL methylation assays.

### *In vitro* kinase assays (Haspin)

Kinase assays were performed using the ADP-Glo Kinase Assay kit (*Promega*) as follows. Haspin is reported to auto-phosphorylate *in vitro* [41–43]. Optimal enzyme concentration for endpoint analysis was identified through an enzyme titration (0-500 nM) against 1 µg of chicken erythrocyte oligonucleosomes according to the procedure described below (**Figure S4F**). To improve the ratio of signal from phosphorylation of the nucleosome to signal of Haspin autophosphorylation in our assays, we performed an additional pre-autophosphorylation step. Purified Haspin (465-498; 250 nM) was incubated for 1 hour at room temperature following addition of 1 mM ATP (*Promega*) in a reaction volume of 50 µL (in 50 mM Tris-Cl pH 7.6, 5 mM MgCl_2_, 1 mM DTT, 0.01% Triton X-100). Following incubation, the autophosphorylation mix was incubated with 25 µL Ni-NTA agarose beads (pre-equilibrated in reaction buffer) for 15 minutes at 4° C. Beads were pelleted by brief centrifugation and supernatant (unbound) was removed. Beads were then washed three times with wash buffer (50 mM Tris-Cl pH 7.6, 5 mM MgCl_2_, 1 mM DTT, 0.01% Triton X-100) to remove remaining ATP/ADP. Autophosphorylated Haspin was then eluted in a final volume of 50 µL in elution buffer (wash buffer supplemented with 200 mM imidazole), and fractions were assessed for residual ADP content by ADP-Glo Kinase Assay detection. Autophosphorylated Haspin (25 nM) was then incubated with nucleosome substrate (300 nM) for 30 minutes at room temperature following addition of 500 µM ATP (*Promega*) in a reaction volume of 20 µL (in 50 mM Tris-Cl pH 7.6, 5 mM MgCl_2_, 1 mM DTT, 0.01% Triton X-100). Following incubation, reactions were quenched and analyzed for phosphorylation according to the standard kit procedures.

### Mass Spectrometry Data Analysis

Histone H3 proteoform data from HEK293 (ATCC CRL-1573) ± HDAC inhibition (5 mM sodium butyrate for 2h) were re-analyzed from MassIVE Dataset MSV000091578 [9].

The contribution of site-specific Histone H3 acetylation to H3K4me1 (Figure 1B) was quantified as in Equation 1.

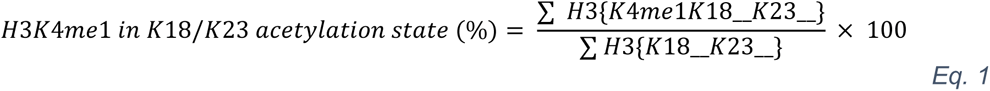

Where ‘___’ denotes K18/K23 acetyl occupancy (un or ac). For example, to determine the contribution of K18acK23ac to H3K4me1 we calculate it as in Equation 2.

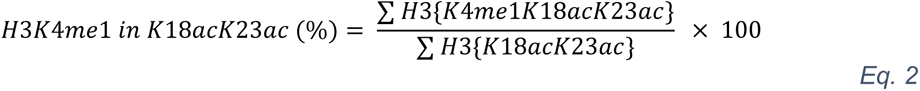

The contribution of site-specific Histone H3 acetylation to H3K4me3 (Figure 2B) was quantified as in Equation 3.

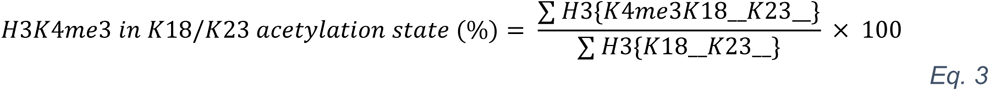

Global/discrete methylation or acetylation as a percentage of total H3.1 (Supplementary Figures S2 & S4) was quantified as in Equation 4.

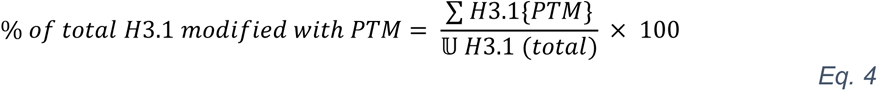

Where ‘𝕌 H3.1 (total)’ denotes the sum of all H3.1 proteoforms. For example, to determine the percentage of H3.1 modified with K4me1 we calculate as in Equation 5.

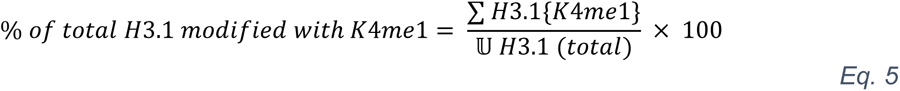

Total H3 N-terminal tail methylation (Supplementary Figure S3) was quantified as in Equation 6.

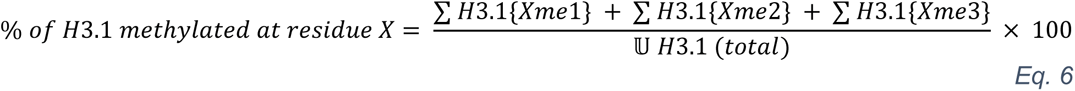

For example, the total H3.1 methylation at K4 was quantified as in Equation 7.

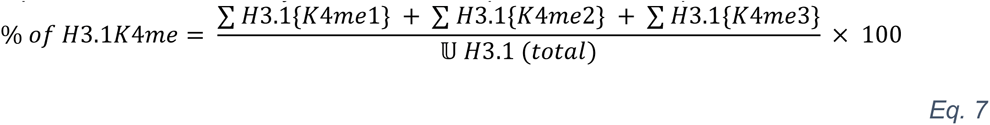

## Data Availability

All data generated for this study is contained within the manuscript and Supporting Information. Previously collected and reported mass spectrometry data that was analyzed in this study is publicly available as indicated in Experimental Procedures.

## Supporting Information

This article contains supporting information.

## Acknowledgments

We thank colleagues for the generous supply of materials (see Experimental Procedures) and members of the Strahl, Young, and Dowen labs for helpful discussions and suggestions. We also thank Michael-Christopher Keogh at *EpiCypher, Inc.* for providing helpful comments on the manuscript.

## Author Contributions

GCF: data curation, formal analysis, investigation, visualization, methodology, writing-original draft, writing-review & editing; KFP: data curation, formal analysis, investigation, methodology, writing-review & editing; BRS and LNV: data curation; JMD: writing-review & editing, funding acquisition; BDS: Conceptualization, formal analysis, funding acquisition, investigation, project administration, supervision, writing-original draft, writing-review & editing; NLY: Conceptualization, formal analysis, funding acquisition, investigation, project administration, supervision, writing-review & editing; KJ: Conceptualization, formal analysis, investigation, project administration, supervision, funding acquisition, writing-original draft, writing-review & editing.

## Funding & Additional Information

GCF is supported by a Predoctoral Training Grant from the National Institute of General Medical Sciences (NIGMS; T32GM135128). KJ is supported by a Postdoctoral Training Fellowship from the National Institutes of Health (NIH; T32CA217824) to the UNC Lineberger Cancer Center and a Postdoctoral Fellowship from the American Cancer Society (PF-20-149-01-DMC). This work was also supported by NIH grants to NLY (R01GM139295, P01AG066606, 1R01AG074540, R56HG012206, R01CA276663 and R01CA193235) and to BDS (R35GM126900).

## Conflicts of Interest

*EpiCypher* is a commercial developer and supplier of fully defined semi-synthetic nucleosomes as used in this study. NNR, BG and BDS own shares in *EpiCypher* with BDS also a board member of same.

---

**Supplementary Figure S1.**
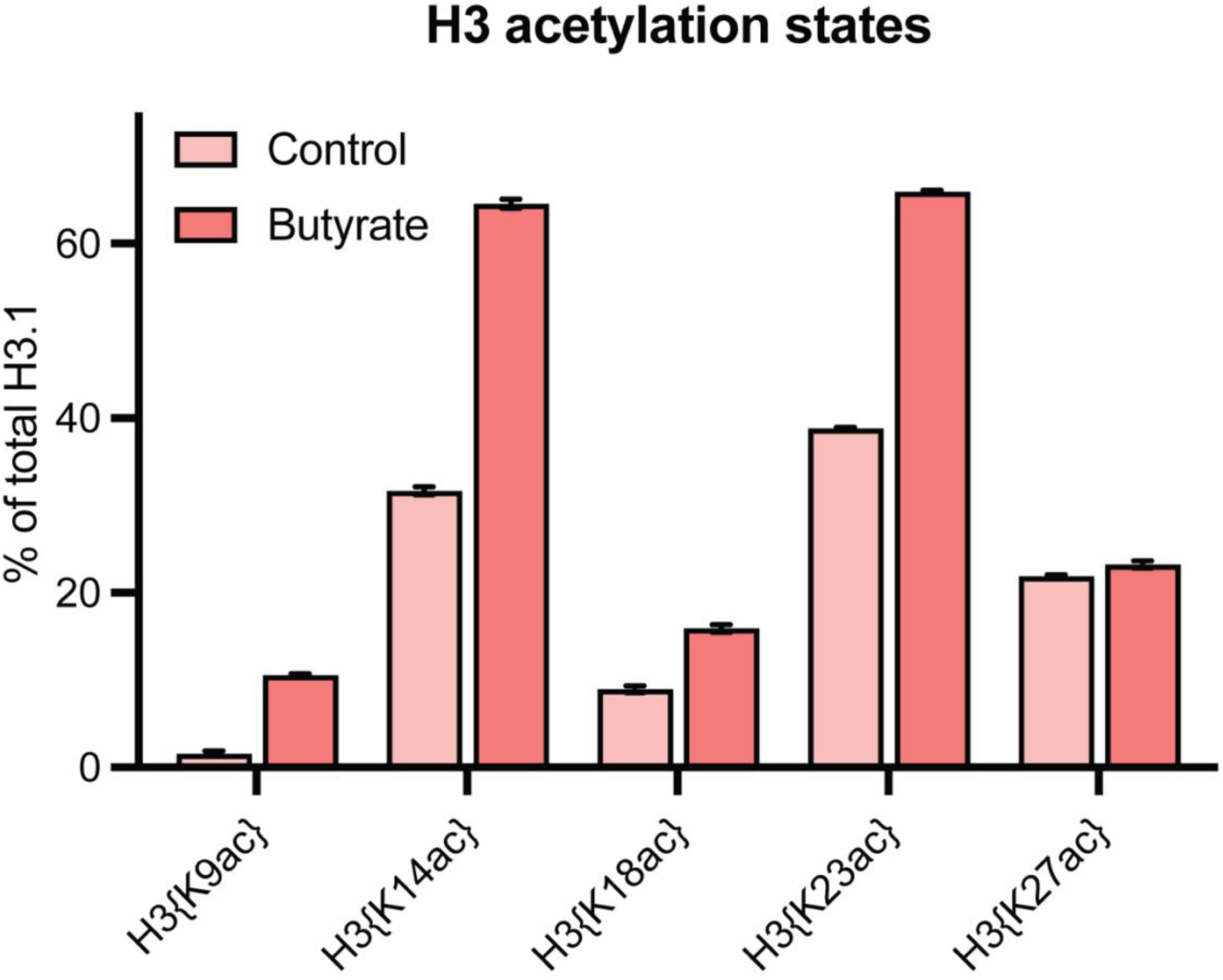
Quantification of total H3 N-terminal tail acetylation. Acetylation states with (dark bars) and without (light bars) HDACi (butyrate) were quantified by middle-down mass spectrometry in HEK293 cells. MS data shown is reported in Supplementary Table 1. Error: SEM.

**Supplementary Figure S2.**
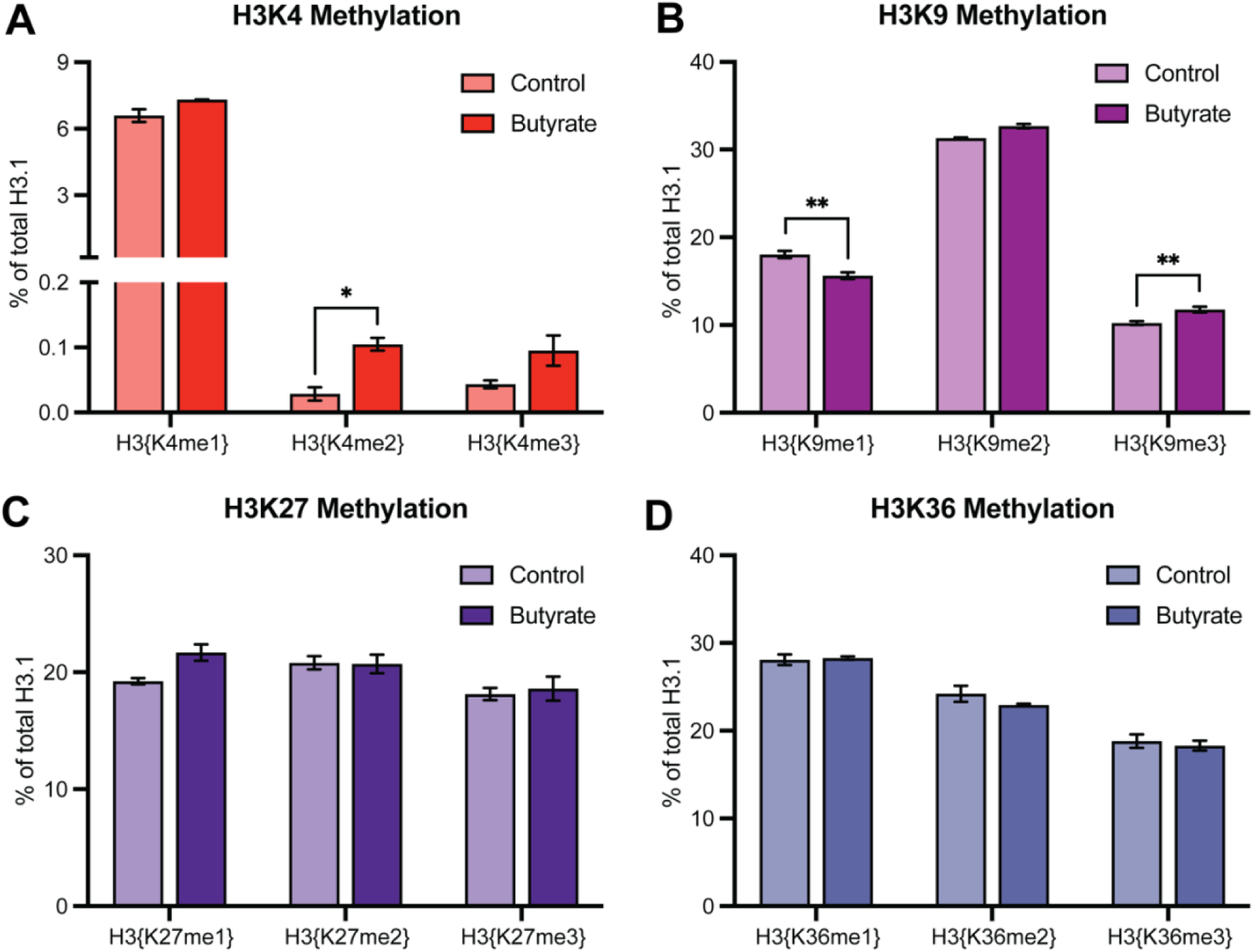
Quantification of global methylation levels with and without HDAC inhibition. H3K4 (A), H3K9 (B), H3K27 (C), & H3K36 (D) methylation states with (dark bars) and without (light bars) HDACi (butyrate) were quantified by middle-down mass spectrometry in HEK293 cells. MS data shown is reported in Supplementary Table 1. Significance was determined by Student’s t-test. NS unless otherwise designated. \**p* < 0.05, \*\**p* < 0.005. *n* = 3. Error: SEM.

**Supplementary Figure S3.**
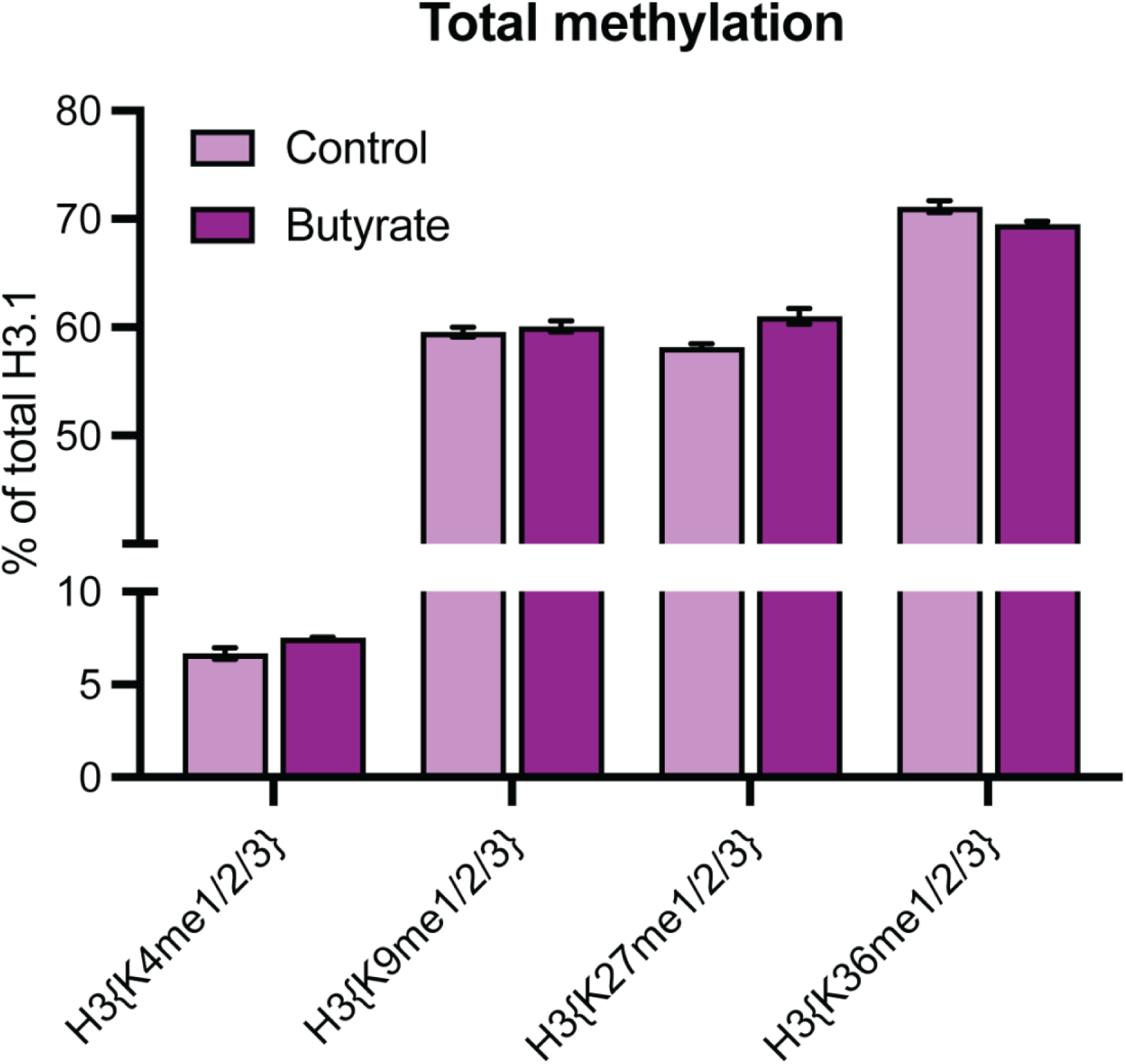
Quantification of total H3 N-terminal tail methylation. Total methylation with (dark bars) and without (light bars) HDACi (butyrate) was quantified by middle-down mass spectrometry in HEK293 cells. MS data shown is reported in Supplementary Table 1. Significance was determined by Student’s t-test. NS unless otherwise designated. Error: SEM.

**Supplementary Figure S4.**
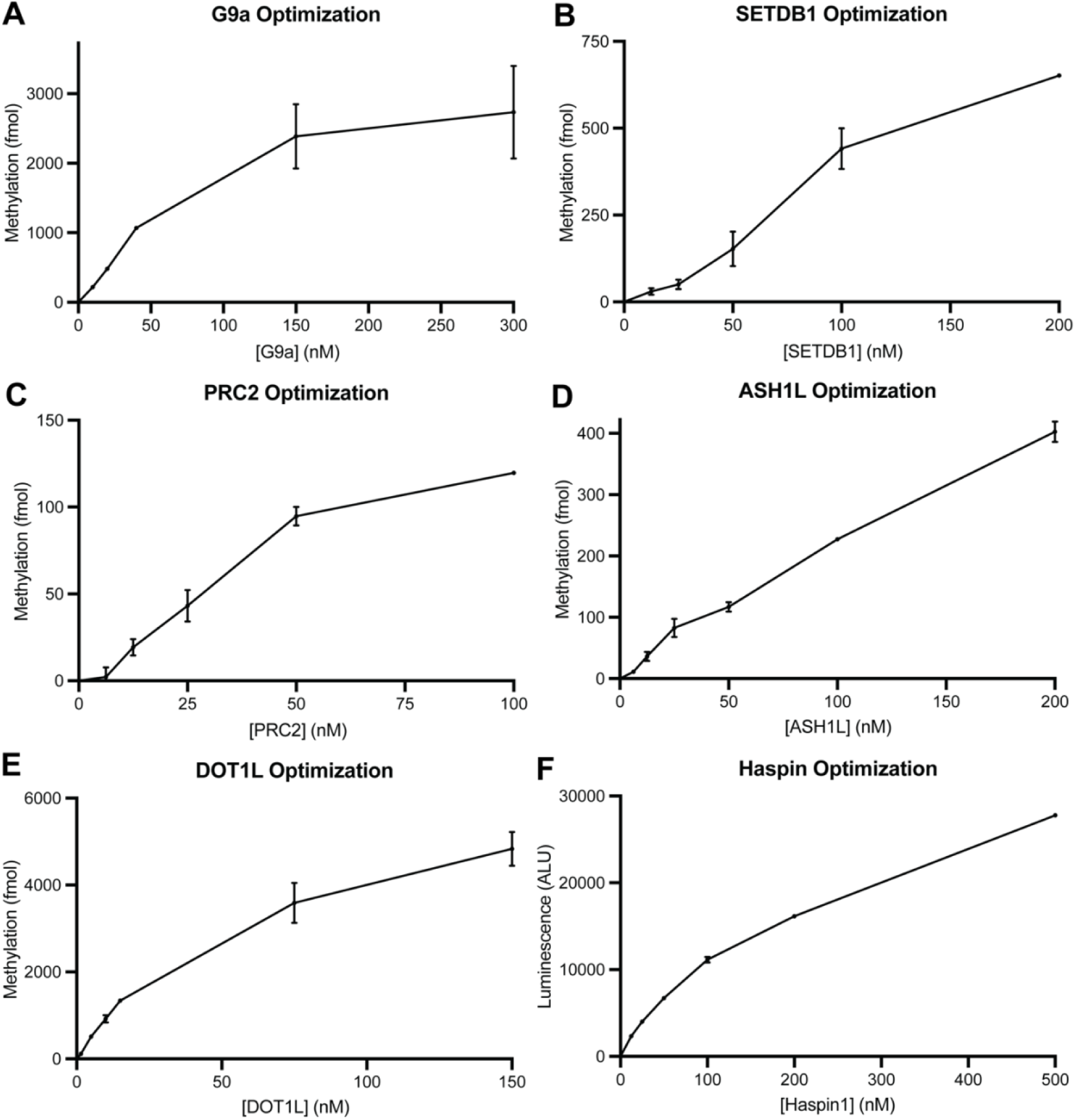
Determination of optimal enzyme concentrations for *in vitro* methylation and kinase assays. Enzymes were titrated against 1 µg of chicken erythrocyte oligonucleosomes. Downstream assessment of either methylation or phosphorylation w as performed as described (see Experimental Procedures). *n* = 2 for all assays shown. Error: SEM.

